# A metapopulation framework integrating landscape heterogeneity to model an airborne plant pathogen: the case of brown rot of peach in France

**DOI:** 10.1101/2023.10.06.561213

**Authors:** Andrea Radici, Davide Martinetti, Chiara Vanalli, Nik J. Cunniffe, Daniele Bevacqua

## Abstract

Plant disease dynamics are driven by the concurrent interplay of host susceptibility, pathogen presence, and environmental conditions. While host susceptibility and local environmental conditions can readily be characterised, the transmission of an airborne pathogen depends on the biotic and abiotic conditions of the surrounding environment.

Here, we propose an original metapopulation framework integrating landscape heterogeneity, in terms of climate and host density, where local populations of plant hosts are connected via air-masses which allow pathogen dispersal. We explicitly account for climatic drivers affecting pathogen release and survival while modelling aerial dispersal using Lagrangian simulations, as well as host phenology and infection. We calibrate the model parameters according to the literature and using Approximate Bayesian Computation against observations of brown rot incidence in French peach orchards from 2001-2020 across an area of 50,000 *km*^2^. We used the model to produce maps of risk, distinguishing site dangerousness (risk of causing secondary infection in other sites) and vulnerability (risk of becoming infected) across the our study area.

We find that most dangerous and vulnerable sites are located along the Rhône Valley, due to the concurrence of high density of peach cultivation, a suitable climate and persistent airborne connections. Our work represents a first step to integrate metapopulation theory, epidemiology and air-mass movements to inform plant protection strategies, and could be adapted to optimize crop protection under future climate projections.

## 1 Introduction

Plant pathogens are a critical issue endangering global food security (Ristaino et al., 2021). Our limited comprehension of long distance dispersed pathogens, *i*.*e*. transported by wind or other vectors over regional to continental scales (Aylor, 2003; Brown and Hovmøller, 2002), has direct consequences on the implementation of plant protection strategies (Cunniffe et al., 2015; Hyatt-Twynam et al., 2017; Parnell et al., 2017). Given the difficulty to eradicate such pathogens, management strategies should focus on preventing emergence via surveillance (Mastin et al., 2020). The appearance of *Monilinia fructicola* in Europe represents an example of an airborne pathogen whose introduction has evaded conventional crop defense measures (EPPO, 2023).

*Monilinia spp*. are fungal species threatening stone fruit production (Bryde and Willets, 1977; Hrustić et al., 2012). *M. fructicola* is an alien species to Europe, initially observed in France in 2001 (Lichou et al., 2002). Despite being classified as quarantine pathogen, in the following years this new strain was progressively observed in central and southern Europe (Oliveira Lino et al., 2016). Such uncontained invasion may be explained considering the efficacy of its dispersal mechanism. The capability of spores to resist UV radiation (Vilanova et al., 2021) and the compatibility of its aerodynamic diameter (Yamamoto et al., 2014) with airborne transport (Wang et al., 2021) combine to suggest that *Monilinia* spores may spread via air-masses (Bryde and Willets, 1977).

Although epidemiological models already exist to study the local dynamics of brown rot (Bevacqua et al., 2018, 2023), its airborne spread remains unexplored. A possible framework to describe brown rot spread at landscape scale consists in coupling in-site epidemiological dynamics and between-sites pathogen dispersal to create a network of spatially distributed hosts connected via a moving pathogen - a metapopulation. This approach has been widely explored in epidemiology (Keeling and Gilligan, 2000; Thrall and Burdon, 2003) with the general assumption of an isotropic spread (Cunniffe et al., 2016; Rimbaud et al., 2018; Mastin et al., 2020; Fabre et al., 2021). However, in the case of airborne plant diseases, scientific research has recently been extended to include realistic patterns of dispersal thanks to the development of Lagrangian models (Draxler and Hess, 1998; Jones et al., 2007). Studies based on such models have explored the airborne dispersal of plant diseases (Sutrave et al., 2012; Meyer et al., 2017), with management implications (Allen-Sader et al., 2019). Nevertheless, these applications have focused largely on transport between known source and sink locations, with no consideration of the coupling of repeated cycles of dispersal, infection and onward spread that characterises epidemics, which could be embedded in a metapopulation framework. Such a framework would help to conceptually disentangle the local factors (climate) from the connectivity (transport of spores) to understand their relative importance to the emergence of the disease.

In this study, we present a model which integrates current knowledge about climate-dependent fruit phenology (Vanalli et al., 2021) and epidemiology (Bevacqua et al., 2018, 2023) where pathogen dispersal among units is described by Lagrangian simulations of air-mass movements to reproduce the disease dynamics over multiple growing seasons at regional scale. We use brown rot of peach in continental France as a case study. We calibrate the model against observations of disease incidence in the last two decades and we verify the importance of including directional airborne transport by comparing the performances against a null model, in which connectivity is modelled with an isotropic dispersal kernel. We use the metapopulation model to produce maps of epidemiological risk, to find where the most endangered locations for infection are located, and where, if a *Monilinia*-like pathogen were introduced, regional disease size would be maximised. These maps represent one possible application of how this model, based on simulated air-mass movements, can feed into management decisions for plant protection at the national scale.

## 2 Materials and methods

### 2.1 Model overview

The geographic domain corresponds to the Safran grid (Bertuzzi and Clastre, 2022), made of square cells 0.11^*°*^ *×* 0.11^*°*^ (∼ 8 *×* 8 *km*^2^, hereinafter referred to as “unit”) overlaid on metropolitain France. We considered those 755 units covered by an important peach orchard area (*>* 0.01 *ha/km*^2^; hereafter “cultivated area”) and not isolated (Fig. 1a; see Section SI1.1).

**Figure 1:**
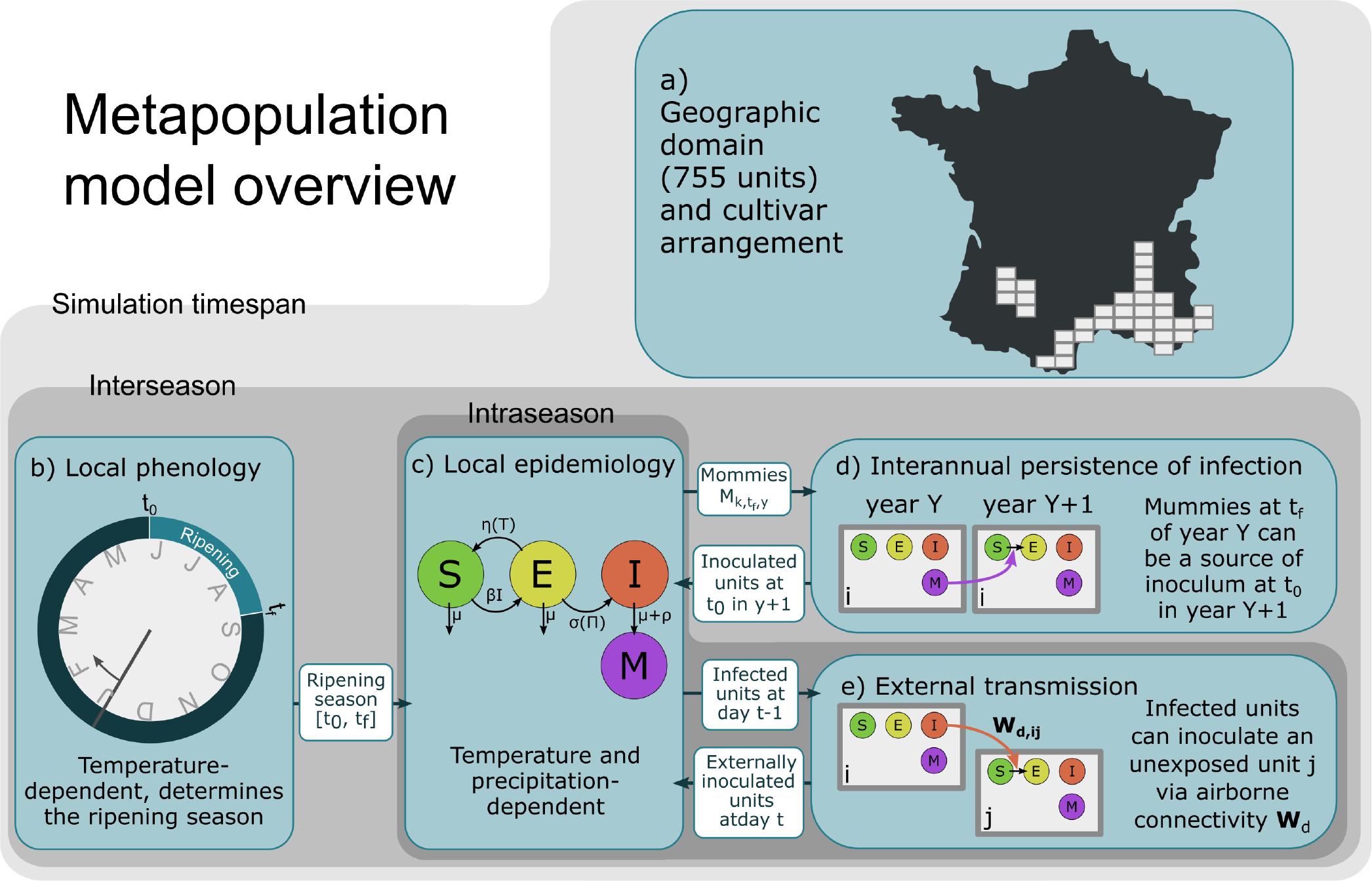
Model overview. a) Geographic domain, composed of 755 ∼ 8 *×* 8 *km*^2^ units in France, assigned with a single cultivar for the whole duration of a simulation; in each unit, b) a climate-dependent phenological model is used to estimate the occurrence of the beginning of ripening *t*_0_ (“pit hardening”) and “end of ripening” *t*_*f*_ (harvest); c) a climate-dependent compartmental model describes the disease dynamics; d) at *t*_0_ of a new season, previous year unharvested mummies determines local inoculum; e) unexposed units can be inoculated at a daily frequency by airborne spores released by infected units.

For every unit, for every year in 1996-2020 we computed the ripening period, from pit hardening *t*_0_ to harvest *t*_*f*_ (Fig. 1b), via a phenological temperature dependant model (see Vanalli et al., 2021 for details), which corresponds to the period of susceptibility. Note that *t*_*f*_ differs among different peach cultivars (i.e. early, mid-early, mid-late, late).

For each unit *i*, we run a type SEI climate driven epidemiological model (see Bevacqua et al., 2023 for details) from *t*_0,*i*_ to *t*_*f,i*_ where *I*(*t*_0_) = 0 and *S*(*t*_0_) +*E*(*t*_0_) = 15 *fruits/m*^2^ (Fig. 1c). Every year, for each unit, we stochastically assess the value of *E*(*t*_0_) as a function of disease incidence in the precedent year. If *E*_*i*_(*t*_0_) *>* 0, the unit is considered as “exposed” and the epidemic dynamics is independent from the epidemic state of the other units. Contrariwise, if *E*_*i*_(*t*_0_) = 0, the epidemic cannot spread unless the inoculum comes from connected infected units. In this case, we calculate the probability that it assumes a positive value *E*_*i*_(*t*) *>* 0 with daily frequency (Fig. 1d). Such probability depends on the incidence of units that act as a source of airborne inoculum (Fig. 1c).

Every year, for each unit, we use the simulated epidemic trajectory to compute the local latent infection, represented by fruits that, following infection, mummify (*M*). We use such a value to assess the probability that, in that unit, a fraction of fruit will be exposed to brown rot at the beginning of the following ripening season (Fig. 1e).

### 2.2 Model Equations

#### Climate dependent phenology and in-unit epidemiological dynamics

We derived initial *t*_0_ and final *t*_*f*_ fruit ripening time according to Vanalli et al. (2021) (Fig. 1b).

We adapted the SEI model for brown rot of peaches developed by Bevacqua et al. (2023) (Eq. 1) to deterministically describe the in-unit disease dynamics, with the additional class *M* (Fig. 1c).

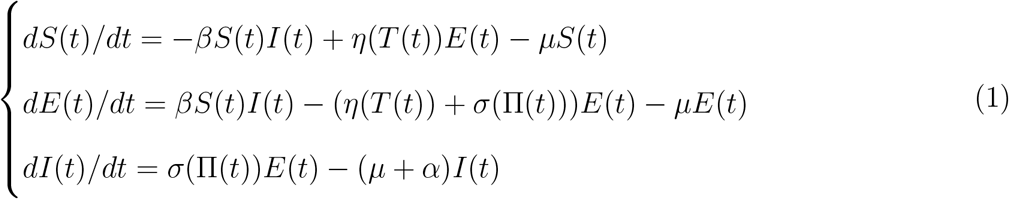

Where *β* is the transmission rate, *η*(*T* (*t*)) is the temperature-dependent spore mortality rate, *σ*(Π(*t*)) is the rain-dependent infection rate, *μ* and *α* are the natural and infection-related abscission rates. The value of *M* is computed at *t*_*f*_ :

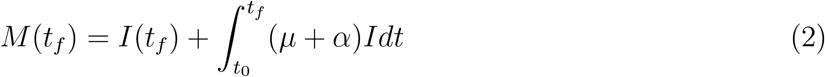

where we include remaining infected fruits at harvest *I*(*t*_*f*_) among the mummies. We used the climate reanalysis provided by Siclima (Delannoy et al., 2022; Caubel et al., 2015).

#### Cultivar distribution across the domain

We inferred the geographical distribution of the cultivars by computing the yield *Y*_*i,z*_ by unit *i* in 1991-2005 in absence of disease, for every cultivar *z*:

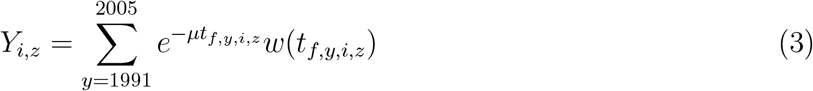

where *t*_*f,y,i,z*_ is the harvest date of variety *z* in unit *i* in year *y*, and *w*(*t*_*f,y,i,z*_) is the relative fruit weight (Eq. SI2). We assumed that the probability *P*_*i,z*_ of presence of cultivar *z* in *i* is proportional to *Y*_*i,z*_ (Fig. SI1):

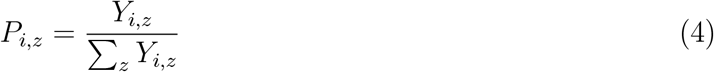

Finally, we created 100 randomized geographical distributions of cultivars, where a unit is associated only to one cultivar, extracted with probability *P*_*i,z*_, for the whole duration of a simulation.

#### Interannual persistence of inoculum

At *t*_0_ some units may have some inoculum which originated by overwintering mummies or other sources (Oliveira Lino et al., 2016; Fig. 1e). We assumed the presence of inoculum as a fraction of the fruit load being already “exposed”: *E*(*t*_0_) = 0.27 *fruits/m*^2^. We determined the probability of having a primary inoculum *P*_*o*_ in year *y* in unit *i* as the complementary of the product of the independent probabilities of not having any inoculum:

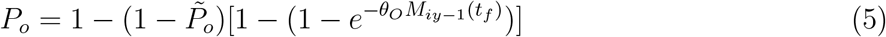

Where 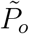, fixed to 0.2 to avoid non-identification issues (see Section SI1.7), represents the probability of having an inoculum due to external sources, while 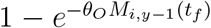 expresses the probability of overwintering. The density of mummies at *t*_*f*_ of the precedent year *M*_*i,y*−1_(*t*_*f*_) is weighted by parameter *θ*_*O*_, to be estimated.

#### External transmission of the inoculum

If instead the unit is unexposed at *t*_0_, an epidemic may be triggered by stochastic introduction of inoculum (Fig. 1e) from infected units. We modelled such inoculation as a Bernoulli variable, with probability *P*_*e*_, depending on the rate of external infection *R*_*i,t*+1_:

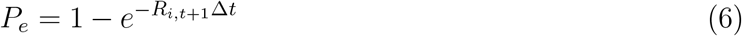

We extracted new daily inoculated units via Eq. 6 (∆*t* = 1*d*). Rate *R*_*i,t*+1_ is defined as:

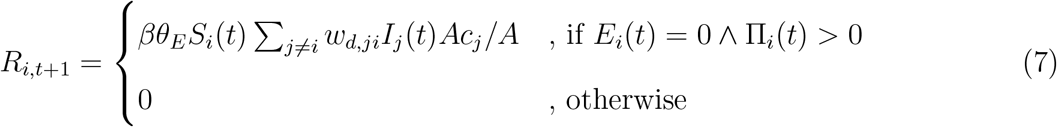

where *β* is the transmission term weighted by *θ*_*E*_, to be estimated, *S*_*i*_(*t*) and *I*_*j*_(*t*) are susceptible and infected fruit loads in units *i* and *j*, term *Ac*_*i*_*/A* corrects to consider units of different cultivated areas, *w*_*d,ji*_ is the airborne transport probability, Π_*i*_(*t*) is the precipitation in the arrival unit *i*, allowing wet deposition.

An inoculation is imagined as a a fraction 0.27 *fruits/m*^2^ of susceptible load that become exposed, as in the case of overwintering. The term *E*_*i*_(*t*) = 0 prevents re-introduction of inoculum.

#### Airborne epidemic network

To estimate airborne transport probability *w*_*d,ji*_ we set up airmass simulations with the HYSPLIT model (Draxler and Hess, 1998). We ran forward Lagrangian trajectory simulations from each unit centroid, 1 m above terrain height, 4 times a day, uniformly extracted from daylight hours (since sunlight facilitates atmospheric turbulence and spore escape; Levetin, 2015) in 2008-2019. We set the travel duration to 6 hours, a compromise between the viability of thin-walled spores (of the order of magnitude of one hour; Oneto et al., 2020) and of thick-walled spores (few days; Visser et al., 2019).

Along a trajectory *v*, we computed the probability of spore survival to temperature and sunlight-induced mortality. We intersected *v* with the domain grid, obtaining a weighted and directed connectivity matrix *W*_*t*_ in which an element *w*_*t,ij*_ represents the probability *P*_*t*_ that *v* started at time *t* in unit *i* crosses unit *j*. We eventually post-processed matrices *W*_*t*_ into *W*_*d*_ by averaging over each day of the year *d* and including an additional connectivity for neighbouring units (see Section SI1.6).

#### Observations of the disease

We collected observations of the disease incidence (as “weak” or “strong”) in different locations from *i*) scientific articles, *ii*) plant health bulletins, *iii*) master thesis reports and *iv*) expert judgement of experimental fields (Radici, 2023).

To compare model outputs with the categorical observations, we mimicked the process by which experts, after examining losses, state whether the incidence has been “weak” or “strong”. Losses *L* are expressed as:

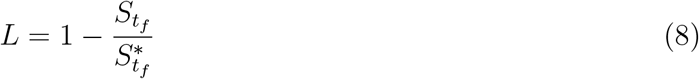

Where 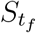and 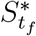 are the actual and disease-free fruit loads at harvest time. We assumed there exists a threshold *θ*_*L*_, to be estimated, which distinguishes high and low losses:

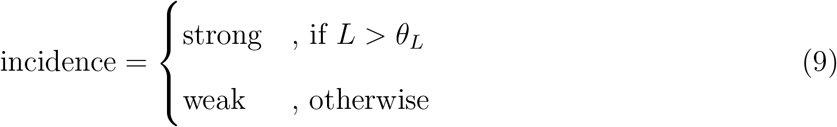

### 2.3 Training and stratified cross validation of the model’s parameters

To estimate the parameter set ***θ*** = (*θ*_*E*_, *θ*_*O*_, *θ*_*L*_) we followed an Approximate Bayesian Computation procedure (ABC; Csilléry et al., 2010; Minter and Retkute, 2019). This allows to estimate the posterior distribution of the parameters by running the model several times and selecting those sets whose performances satisfy a proximity threshold *ϵ* to the observations (see Section SI1.7). We chose Cohen’s *κ* index (Fielding and Bell, 1997) as a measure of proximity (ranging from -1, lowest, to 1); we fixed a value (0.475) and set *ϵ* as 1 − *κ* = 0.525.

To further restrict the parameter sets, we used a Stratified K-Fold Cross Validation algorithm (Arlot and Celisse, 2010). We assessed *κ* both on training (*κ*_*ϕ*_) and on testing (*κ*_*ψ*_) sets, and took the first 100 more frequent accepted sets as our final ensemble ***θ***^**∗**^ (see Section SI1.8).

### 2.4 Testing the null model

We tested the hypothesis of directional airborne epidemic spread against a null model where we replaced wind-driven matrices **W**_**d**_ with an isotropic kernel (matrix **U**). Each element *u*_*ij*_ depends exclusively on the distance between *i* and *j*. We run the two models and we compared their performances through a Monte Carlo analysis (Gotelli et al., 2004; see Section SI1.9).

### 2.5 Estimating risk of brown rot of peaches

We used the model to find the most dangerous units, intended as the ones which, if infected, would maximize the disease size and the most vulnerable units, intended as the ones which become easily infected because of an outbreak elsewhere in the domain. First, we set a unit to be completely infected; then, we associated a random year in 1991-2010, a randomized geographical rearrangement of cultivars, a random parameter set from the posterior distribution, and we ran a 10-years simulation, computing losses in the last simulated year according to Eq. 8.

In this experiment we set the parameter 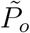 (external inoculum) to 0 until that unit has been infected (Eq. 5). From 75,500 simulated epidemics (755 units *×* 100 stochastic repetitions for each unit) we computed two indices: *i*) the “vulnerability”, i.e. the average local losses by secondary infection, started anywhere in the region; *ii*) the “dangerousness”, i.e. the overall average losses (weighted by cultivated area) caused by an infection in that unit (see Section SI1.10).

## 3 Results

### 3.1 Wind connectivity matrices

Air-mass connectivity, summarized by the annual matrix **W** (the weighted average of all the daily matrices **W**_**d**_; see Section SI1.6), varies heterogeneously through the study area and can be represented as a spatial network (Fig. 2a, rearranged on regions defined in panel b). Beside self loops, an important spread pathway occurs in the Rhône Valley (regions D-E-F; Fig. 2b) followed by a Mediterranean coastal pathway (regions B-C-F-G-H). On the opposite, the northeastern part of the domain (A, Guyenne) has very few connections with the rest of the domain.

**Figure 2:**
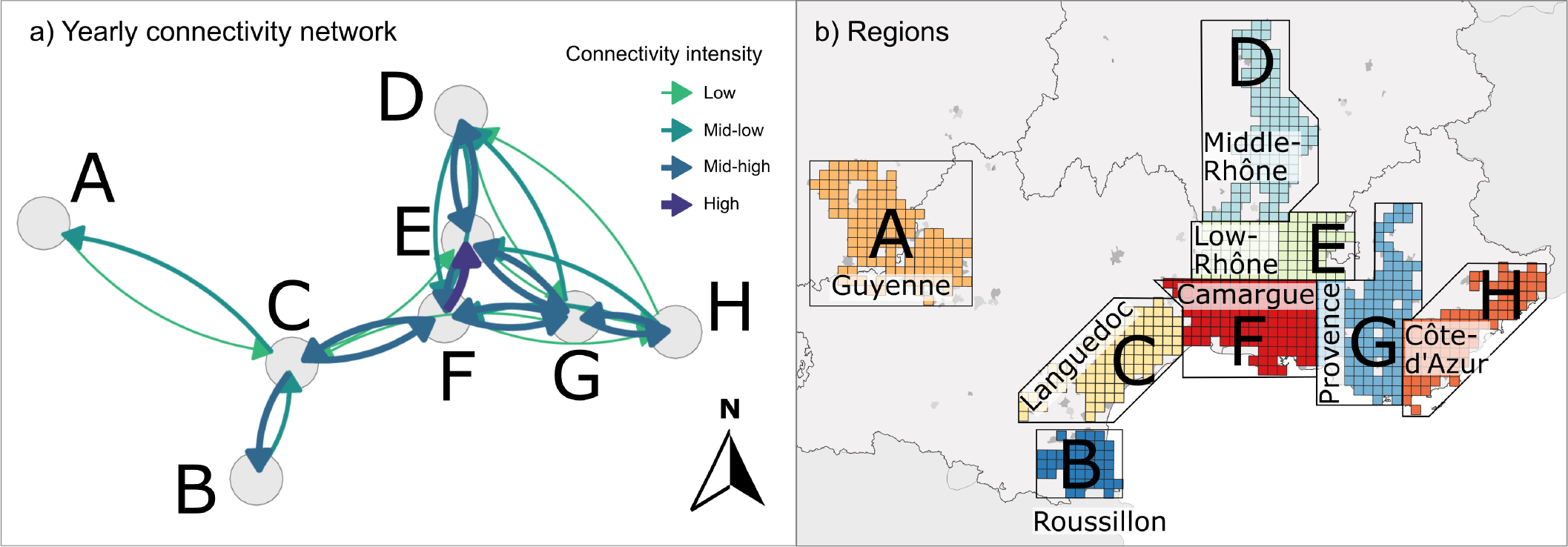
Average connectivity matrix *W* a) as an oriented network, rearranged grouping units into b) eight regions. The colour scale represents connectivity on a logarithmic scale (High: probability of a daily deposition of spore from any unit in a region to any of a second region in the order of 10^−4^; Mid-high, 10^−5^; Mid-low, 10^−5^; Low, 10^−7^) consistently with Fig. SI2.

### 3.2 Parameters estimation

Depending on the cross validation splitting, the size of the accepted parameter sets varies from 7 to 67 (average = 29) over 200,000. Cohen’s (*κ*_*ϕ*_) assessed on the training set has an average value of 0.5 (min = 0.496; max = 0.502), while on the testing set (*κ*_*ψ*_) has an average value of 0.2 (0.046, 0.409).

The losses threshold over which an epidemic is considered as “strong”, represented by parameter *θ*_*L*_, has average value is 30.3% (interquartile range: 27.1% to 32.9%; Fig. SI3c). It is more complicated to directly visualise the range of accepted values of *θ*_*E*_ (the weighting parameter driving external inoculum; average value = 1.8 *×* 10^−6^; 6.1 *×* 10^−7^ to 2.4 *×* 10^−6^; Fig. SI3b) and *θ*_*O*_ (the weighting parameter driving overwintering; average value = 1.0 *×* 10^−2^; 7.1 *×* 10^−3^ to 1.4 *×* 10^−2^, Fig. SI3a). We therefore computed the “average annual number of external infections”, which is function of *θ*_*E*_ (average value = 106.2 out of 755; 48 to 148; Fig. SI4b) and the “average number of units infected at *t*_0_” which is function of *θ*_*O*_ (average value = 159.7 out of 755; 149 to 168; Fig. SI4b).

The average value of *κ* computed on the whole observations is 0.41 and the correct classification rate (the ratio of true positive and true negative, Fielding and Bell, 1997) is 71.8%. This latter varies throughout each cultivar (Fig. 3; Fig. SI5a). It is generally higher for “weak” incidences (79.7%) and lower for “strong” incidences (61.1%). Also, it decreases passing from early (78.4%) to late cultivars (62.7%; Fig. 3, SI5c).

**Figure 3:**
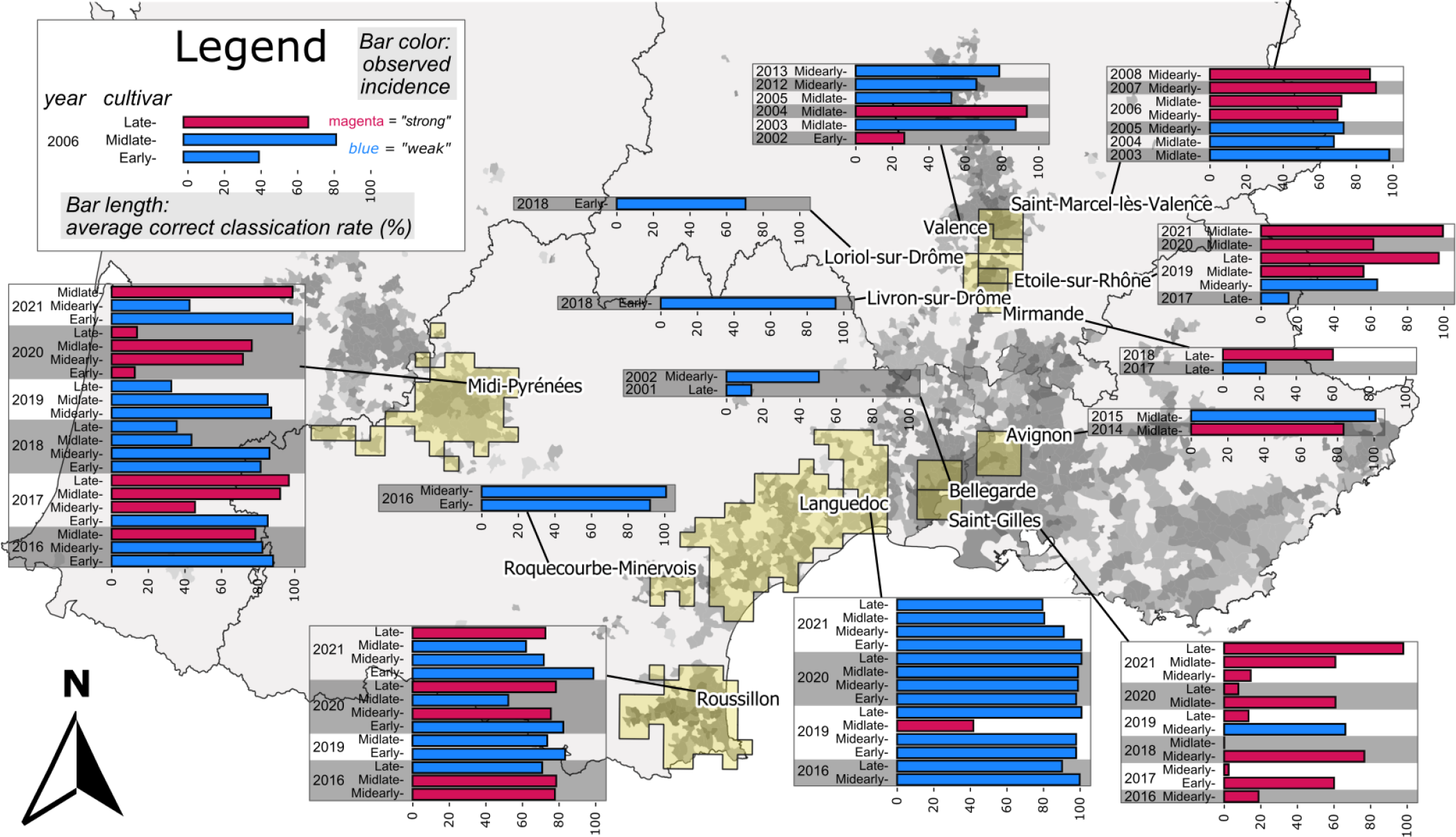
Observations of incidence of brown rot in peach orchards and estimated model accuracy. The observed incidence is represented by the color. Locations are mapped as yellow shades. The bar length represents the correct classification rate of the model, i.e. the % of true positives and true negatives. Gray shades represent cultivated areas per municipality.

The Monte Carlo analysis revealed that the wind-driven model performs significantly better that the null one (p-value *<* 5 *×* 10^−6^).

### 3.3 Dangerousness and vulnerability to brown rot

The most dangerous units (average dangerousness, i.e. the average regional losses caused by an infection in that unit) values are found in the Camargue (region F in in Fig. 4a, 25.7%, with a maximum of 38.9%), followed by other regions along the Rhône Valley (24%): the Middle-Rhône (D, 23.8%) and Low-Rhône (E, 21.9%). Provence (G, 9.5%), Languedoc (C, 8.7%) and Côte d’Azur (H, 5.9%) display intermediate values, while Roussillon (B, 4.8%,) and Guyenne (A, 2.6%) have to the lowest average values.

**Figure 4:**
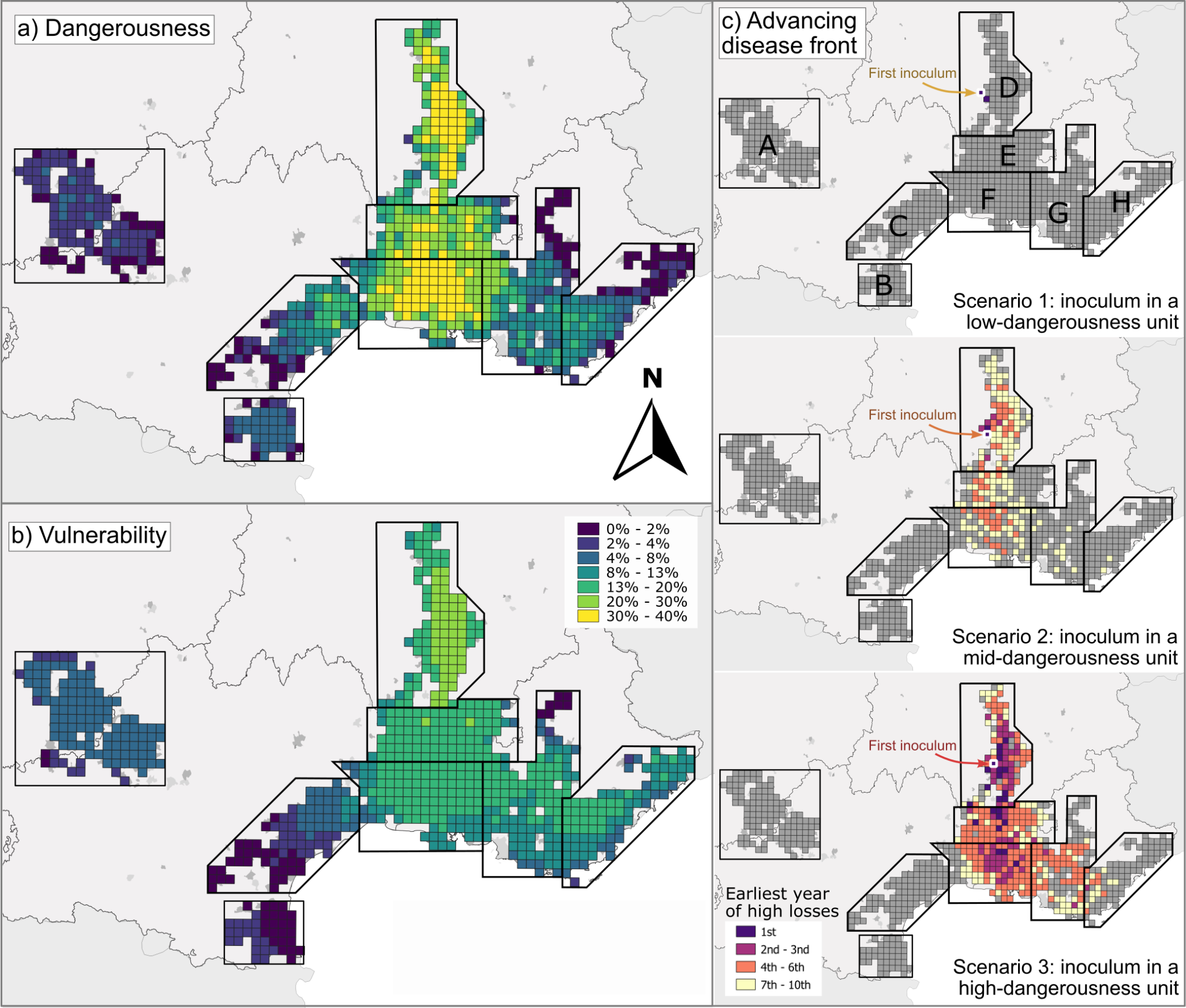
Geographical distribution of epidemiological risk. a) Dangerousness (%), i.e. the average regional losses 10 year after inoculation; b) vulnerability (%), i.e. the average local losses 10 year after inoculation of a random unit; c) earliest year of high losses (*>* 30%) in a rapid-spread simulation (i.e., the simulation corresponding to the 67^th^ quantile of cumulative losses) with a single inoculated unit in Middle Rhône (first inoculated unit has a low, middle and high dangerousness in scenario 1, 2 and 3 respectively).

The most vulnerable units (average vulnerability, i.e. the average local losses by secondary infection, of 19%) are again found in in the Middle-Rhône (Fig. 4b), with peaks of 23.9%. From the most to the least vulnerable units, we found in Low-Rhône (average = 16.9%) and Camargue (average = 14.1%) and than eastwards in Provence (average = 11.6%) and Côte d’Azur (H, 11.1%). Guyenne follows with values of vulnerability around 5.2%. Languedoc and Roussillon have again the lowest average values of 3.7% and 2%.

In view of this result we decided to illustrate typical disease dynamics. We therefore recomputed 200 simulations for a specific units. We chose three geographically close units (in Middle-Rhône) but different in term of dangerousness. For each one, we mapped the progress of the advance front of a “rapid spread” simulation, i.e. the earliest year of high losses (*>* 30%) in the simulation generating the 67^th^ (i.e., 2*/*3) quantile of cumulative losses over the 10 years time span (Fig. 4c).

In 11.5% of the simulations where the disease starts in a high-dangerousness the outbreak dies out before the 10^th^ year. These percentage increases to 60.5% for a mid-dangerousness and 85.5% in a low-dangerousness unit. A high-dangerousness unit can provoke high losses in 352 units out of 755, with a main north-south spread direction, the double of a mid-dangerousness (180 out of 755), while a low-dangerousness unit would provoke high losses in only one other unit.

## 4 Discussion

We presented a metapopulation model explicitly including wind-driven dispersal for studying the spatio-temporal dynamics of airborne plant pathogens and we used it to produce maps of disease risk. Our framework allows to match observations of disease incidence while considering the complementary role of local overwintering and the wind-driven transport of inoculum between orchards (Fig. 3): the best fit is obtained when extinctions due to missed overwintering are compensated by re-introductions due to air-masses movements (Fig.s SI3 and SI4).

The role of directional wind-driven transport is corroborated by the Monte Carlo analysis with the null model, fed with an isotropic kernel. Such kernel is typically assumed in scientific literature in absence of knowledge on pathogen transports (Cunniffe et al., 2016; Rimbaud et al., 2018; Mastin et al., 2020; Fabre et al., 2021). Lagrangian trajectory simulations of air masses have been already used to investigate aerial connectivity (Radici et al., 2022); our experiment suggests that, for those pathosystems where airborne transportation is ascertained, it be efficiently integrated in a dynamical framework, helping describing the epidemic spread more accurately than a isotropic kernel.

Defining an acceptance threshold is usually a critical point in the ABC (Minter and Retkute, 2019), but a rooted scientific literature helps interpreting the meaning of *κ*. Negative values characterise models which are worse then random, values over 0.21 are considered as “fair”, while over 0.41 are “good”, “moderate” (McHugh, 2012; Fielding and Bell, 1997). We set *κ >* 0.475 over each cross-validation repetition. The same sets of parameters, if assessed against the testing set, have values of *κ* close to “fair” (Tab. SI2).

The model predicts “weak” incidences better (average correct classification rate = 80%) than “strong” ones (61%). This is consistent, since a “strong” incidence needs both favourable environmental conditions and inoculum presence, while “weak” incidences simply require one condition to be absent. Geographical distribution of model performances can be discussed under this consideration (Fig.s 3, SI5b); “weak” infection levels in Roquecourbe-Minervois, Livron-sur-Rhône, Livron-sur-Rhône and Linguedoc are better identified than “strong” levels in Saint Gilles and Valence. Disease dynamics in early cultivars are generally better described that in late cultivars (Fig.s 3, SI5c), which have by definition longer ripening duration in which they may be infected.

### The risk of brown rot in France

The model identifies both the most dangerous disease spreaders and most vulnerable pathogen receivers along the Rhône Valley (regions D-E-F).

From an management point of view, vulnerability - which can be compared to the secondary infection risk by Meentemeyer et al. (2011) - calls for direct protection (e.g., via fungicide application) of an epidemic unit. Dangerousness, instead, measures the impact of an outbreak at a broader scale; it underlies a vision in which local management has repercussion all over an inter-connected landscape. Independently of its vulnerability, preventing disease in a dangerous unit may be globally advantageous over a larger share of the domain.

Despite being two aspects of the main phenomenon, vulnerability and dangerousness are statistically distributed differently through the domain. The coefficient of variation for the former (0.56) is lower compared to that for the latter (0.84). This means that units are more easily characterised as having high or low dangerousness, rather than for their vulnerability, as in this case they are clustered around to the average value. This suggest that good disease spreaders are more easily distinguishable from bad ones than good or bad pathogen receivers.

Proposed epidemic risk indices depend indirectly from a number of factors: length of ripening, host surface, favourable environmental conditions and air-mass connections. Host density per unit increases proportionally the capacity of infecting connected units, and consequently dangerousness (Eq. 7). Rain occurrence can be a measure of suitability of the environmental conditions triggering infections, either due to local inoculum or wind-mediated (due to wet deposition). In our model, rain triggers the rate *σ* at which exposed fruit load become infected (Fig. 1c, Eq. 1), proportional also to peach weight. In fact, also plant phytosanitary bulletins warn local producers to pay attention to rain when fruits are close to ripening, as this can facilitate cuticle cracking and subsequent infection (Oliveira Lino et al., 2016).

Climatic factors affecting epidemic risk may be more easily visualized in Guyenne and Roussillon due to their low connectivity to the domain (Fig.s 2a, SI2a). In fact, despite scoring low on both indices, their relative isolation from the rest of the domain helps disentangle the importance of local environmental variables from those influenced by connectivity. Guyenne is characterised by little cultivated area (about 5 ha per unit, Fig. SI1e). On the other hand, in this region we observe the largest frequency of rainy days during ripening (24.5 days on average; Fig. SI1e), which is the main factor driving infection in the SEIM model, which may explain why, despite its isolation, its vulnerability is higher compared to other regions such as Languedoc. Roussillon, instead, hosts the largest cultivated areas (33.5 ha per unit) and is slightly more dangerous than Guyenne.

Regions along Rhône Valley, which are then both dangerous and vulnerable, are interested by the typical North-South wind circulation (the Mistral), whose units host large cultivated areas (from 15 to 29.2 ha per unit) and intermediate values of rainy days (12 to 21 days per season). Among mid-risk, but well-connected regions, late varieties are common in Languedoc and Côte d’Azur, and less in Provence (Fig. SI1e). By contrast, they all share little of cultivated areas (5.1 to 7.9 ha per unit) and infrequent rainy events (11.8 to 14 days per season).

Intuitively, the latest varieties should be also the most vulnerable, provides more time for infection. In contrast, these varieties, which typically grow in warmer climates, are relatively common in lid- and low-risk regions (Fig. SI1d). This suggests that frequency of rainy days (which seems to trigger vulnerability) or host surface and connectivity (which are likely to increase dangerousness) may be more relevant epidemiological factors.

The first European detection of *M. fructicola* occurred in 2001 in the Gard department (Lichou et al., 2002; Fig. SI2b, between regions F and E), which hosts dangerous units (Fig. 4a). While epidemics starting from these units persist for several years, their spread is unlikely to cross national borders (Fig. 4c). This slow progression may be attributed to the hypothesis underlying the fitting of the model: assuming that the disease is already widespread everywhere may have led to an underestimation of its aggressiveness.

Moreover, it may be imagined that wind is not the only dispersal medium. Brown rot of peaches is also known to cause high post-harvest losses (Oliveira Lino et al., 2016) and it can be hypothesized that the handling of returnable crates used to ship and store harvested fruits contributes to the spread of the disease (Bryde and Willets, 1977). If one wanted to integrate this transportation mode to our modeling framework, this would be represented by an additional connectivity layer reflecting storage and trading (Hernandez Nopsa et al., 2015). Furthermore, this additional spread would not necessarily occur simultaneously with the ripening season but later, thereby extending the period during which susceptible fruits can come into contact with the pathogen. Beside additional dispersal medium, the discrepancy between observed and modeled spread leaves open the debate about possible multiple introductions of *M. fructicola* in Europe.

### Towards metapopulation-based plant protection strategies

Fungal plant disease dynamics have been traditionally interpreted only under the light of local environmental conditions (Juroszek et al., 2020), neglecting transport of spores. The presence itself of *Monilinia* viable spores correlates with local environmental variables (Holb, 2008), but this information has not been used so far to retrace epidemiological dynamics going beyond local descriptors.

Airborne transport has gained increasing scientific evidence to be among the fastest way of pathogen transport (Schmale III and Ross, 2015), provided that given biological protections (melanin, leading to spores resisting UV light) and aerodynamic features are ascertained (Levetin, 2015). The growing availability of computationally efficient Lagrangian models to reconstruct airmass trajectories, such as HYSPLIT (Draxler and Hess, 1998), allows more research on airborne plant pathogen spread (Radici et al., 2022).

Our study represents a first step to use air-mass movements to both study airborne disease spread among fruit trees and inform plant protection strategies at the national range. This research supports a shift in the current paradigm regarding the optimal spatial scale of disease management from the field to the landscape (Radici et al., 2023; Thompson et al., 2016). Moreover, it could be adapted to perform optimization of epidemic surveillance and disease control under future climate projections.

## Supporting information

Supplementary methods and results

## Acknowledgement

The authors acknowledge the support of funding from the French National Research Agency (ANR) for the BEYOND project (contract # 20-PCPA-0002) and the SuMCrop Sustainable Management of Crop Health Program of INRAE that supported the work of all authors. AR acknowledges the financial support of the *Perdiguier Programme* scholarship from the University of Avignon which allowed his stay at the University of Cambridge as visiting.

Safran climatic data are provided by Météo-France and were downloaded via the SICLIMA platform developed by AgroClim-INRAE.

The calibration of this possible would not have been possible without the help of the “National working group on *Monilinia*”, coordinated by Bénédicte Quilot-Turion. Part of the analysied data were provided by SudExpé and SEFRA applied research stations and INRAE UERI and GAFL units.

We thank Samuel Soubeyrand, Cindy Morris, Renato Casagrandi, Lorenzo Mari, Leonardo Miele, Paco Melià and Dario Constantinescu for valuable suggestions.

The authors declare no competing interests.

## Data Availability Statement

The code that supports the findings of this study is available at the repository [soon available].

Correspondence and requests for materials should be addressed to Davide Martinetti.

