## Supplementary methods and results for "A metapopulation framework integrating landscape heterogeneity to model an airborne plant pathogen: the case of brown rot of peach in France"

### 1 Supplementary Methods

#### 1.1 Domain extraction

We built the domain upon units extracted from the so-called Safran grid (Bertuzzi and Clastre, 2022), which is composed of regular square cells with spatial resolution of $0.11° \times 0.11°$ ($\sim 8 \times 8$ $km^2$). For each unit, we computed stone fruit orchard area and peach orchard area (this latter is referred simply as "cultivated area" in the main text: here, for sake of clarity, we specify that we mean the peach cultivated area). We extracted stone fruit cultivated areas from data collected by the 2010 national French agricultural survey (RGA) conducted by the French Minsitry of Agriculture (*Ministère de l'Agriculture et de la Souveraineté alimentaire*; RGA (2010)). From this database, we summed "stone fruits" (column G_1013_LI_DIM2 == "fruits à noyau") area data (column "Superficie correspondante (hectares)") of all farms (column "G_1013_LIB_DIM1" == "Ensemble des exploitations (hors pacages collectifs)" by municipality (expessed by the INSEE code).

This operation allowed us to obtain the area $As_m$ covered with stone fruit orchards by municipality $m$.

We computed the stone fruits density by municipality and geographically intersected the corresponding shapefile with Safran grid. This operation allowed to associate each unit of the Safran grid with a stone fruit density. Open source software QGIS 3 (QGIS Development Team, 2022) was used for this operation.

We obtained the spatial data about peach orchards form Eurostat at NUTS2 (Eur, 2021), which correspond to groups of French regions (as defined before the 2016 reform: FR1 *Île de France*, FR2 *Bassin Parisien*, FR3 *Nord-Pas-de-Calais*, FR4 *Est*, FR5 *Ouest*, FR6 *Sud-Ouest*, FR7 *Centre-Est*, FR8 *Mediterranée* and FR9 the *Département d'Outre Mer*). We considered data about "Peach and apricot trees - Area by density classes and group of cultivars (area in ha)".

We grouped together all areas from following categories: "Dessert peach and nectarine trees [PCD]"; "Peaches for fresh consumption [PCD_PEA]"; "Yellow flesh peaches [PCD_PEAY]"; "Very early yellow flesh peaches [PCD_PEAY_VE]"; "Early yellow flesh peaches [PCD_PEAY_E]"; "Medium-early yellow flesh peaches [PCD_PEAY_M]"; "Late yellow flesh peaches [PCD_PEAY_L]"; "White flesh peaches [PCD_PEAW]"; "Very early white flesh peaches [PCD_PEAW_VE]"; "Early white flesh peaches [PCD_PEAW_E]"; "Medium-early white flesh peaches [PCD_PEAW_M]"; "Late white flesh peaches [PCD_PEAW_L]"; "Doughnut peaches [PCD_PEAD]"; "Medium-early yellow flesh nectarines [PCD_NECY_M]"; "Late yellow flesh nectarines [PCD_NECY_L]"; "White flesh nectarines [PCD_NECW]"; "Very early white flesh nectarines [PCD_NECW_VE]"; "Early white flesh nectarines [PCD_NECW_E]"; "Medium-early white flesh nectarines [PCD_NECW_M]"; "Late white flesh nectarines [PCD_NECW_L]"; "Peach and nectarine trees for industrial processing (including group of Pavie) [PCI]".

This operation allowed us to obtain the area $Ac_n$ covered with peach orchards within NUTS2 region $n$. We estimated area $Ac_m$ covered with peach orchards by municipality $m$ located in NUTS2 region $n$ via Eq. 1. This equation assumes that peach cultivated area in municipality $m$ is proportional both *i*) to the stone fruit cultivated area in that municipality $As_m$ and *ii*) to the peach cultivated area in the region where that municipality is located $Ac_n$ and that *iii*) the sum of the peach cultivated area by municipality $m$ is equal to the peach cultivated area in that region $n$.

$$Ac_m = Ac_n \frac{As_m}{\sum_{m \in n} As_m} \tag{1}$$

Each unit of the grid is thus associated with *i*) its area, *ii*) the stone fruits cultivated are and *iii*) the peach cultivated area. Since keeping all the elements of the grid in the domain would imply considerably slower computations describing the dynamic of a great amount of poorly informative units, we decided to keep only those units respecting a threshold of density (i.e., at least 0.01 $ha/km^2$) and continuity (i.e., only units which form groups of at least 4 contiguous elements; Fig. SI1e). By excluding cells with a low density and with no neighbours we discarded a great amount of cells where peach cultivation is known to be absent, for climatic reasons (for example in central and northern France) but Eq. 1 may still assign a residual $Ac$ due to a misclassification error. For instance, we found few, isolated cells in Alsace-Lorraine (north-east of France) where we computed a relevant peach cultivated area. The same cells correspond to a well-known production basin of mirabelle plums, a stone fruit typical of the region and well adapted to its climate. We found more reasonable to conclude that these cells were hosting important mirabelle plum orchards, rather than peach ones; therefore, we proceeded to remove them.

#### 1.2 Pit hardening date

We computed peach growth increase according to equation (Bevacqua et al., 2023):

$$w(t) = \frac{w_B w_M}{w_B + (w_M - w_B)e^{-h(t-t_B)}} \tag{2}$$

Where $w(t)$ is the weight at time $t$, $w_B = 0.49\ g$ is the fresh fruit weight at bloom time, $w_M = 214\ g$ is the fresh maximum fruit weight, $h = 0.056\ (d^{-1})$ is, at first approximation, the conversion rate of resources into fruit mass, $t_B$ is the bloom time.

We used this equation to compute $w_H = 164g$, that is weight at harvest time $t_f$, from 2014 and 2015 data ($t_{f,2014} = 196$, $t_{B,2014} = 59$, $t_{f,2015} = 198$, $t_{B,2014} = 74$, (Vanalli et al., 2021); times are here expressed in terms of day of the year).

As suggested in Bevacqua et al. (2023), once the weight at harvest is known, the conversion rate $h$ should be iteratively recomputed to be applied in other ripening seasons. Therefore, we recomputed $h'$ from the known ripening date $t_R$ for the computation of the pit hardening date ($t_0$):

$$h' = -log(w_B/(w_M - w_B) * (w_M/w_H - 1))/(t_R - t_B) \tag{3}$$

And $t_0$ as follows:

$$t_0 = t_B - log(w_B/(w_M - w_B) * (w_M/w_C - 1))/h' \tag{4}$$

Where $w_C = 61\ g$ is the fresh fruit weight threshold for cuticle cracking.

#### 1.3 Initialization of the SEIM model

In inoculated units, we initialised fruit load as $S_0 = 14.73\ fruits/m^2$, $E_0 = 0.27\ fruits/m^2$, $I = 0\ fruits/m^2$ (to account for the observation according to which, in infected orchards, 1.8% of the fruits are considered as exposed; Bevacqua et al. (2023)) while unexposed units are initialized as $S_0 = 15\ fruits/m^2$, $E = I = 0\ fruits/m^2$. We chose an initial total fruit load of 15 $fruits/m^2$ since this is the average of 2014 and 2015 data in Avignon (Bevacqua et al., 2023).

#### 1.4 Weather variables

We used Safran weather reanalysis data (via the Siclima portal: Delannoy et al. (2022); Caubel et al. (2015)), which are arranged according to the Safran grid (Bertuzzi and Clastre, 2022) the same used for the metacommunity model.

Daily precipitations ("preliq_q, meaning daily liquid precipitations 06-06 UTC"), daily mean temperature ("preliq_q, 01-00 UTC") and max temperature ("tinf_h_q") from the 1st of January 1980 to the the 31st of December 2021 have been stored.

The Safran grid has been recently rearranged: with respect to the original grid, comprising 8602 units, Safran data managers added 379 new units, located on the coasts. This rearrangement created some mismatches with the previous one, since stone fruits have been mapped with the most recent version, while available weather reanalysis are consistent with the previous one. To solve this mismatch, we attributed to the newly introduced units, all located next to coastal environments, the weather series of the neighbouring unit that shares the longest geographical border. The IDs (consistent with the old Safran grid) of the stored variables are indicated in "ID_safran_grid.csv" in supplementary material.

#### 1.5 Determination of the probability of spore deposition $P_v$

A trajectory $v$ as emitted from our HYSPLIT simulations is made up of points in the atmosphere $x_{v,t}, y_{v,t}, z_{v,t}$, with $t = 0, 1, \ldots, 6$, which we linearly interpolate to obtain a continuous trajectory $x_v(t), y_v(t), z_v(t)$. Along a trajectory we assessed spore viability to temperature T (Bevacqua et al., 2023) and other losses, such as solar radiation, deposition and dilution, combined with an exponential kernel, as in eq. 5:

$$P_{v0}(x_v(t), y_v(t), z_v(t), T) = e^{-\alpha t} \int_0^t e^{\frac{-r}{24T(x_v(\tau), y_v(\tau), z_v(\tau))}} d\tau \tag{5}$$

With $r = 1.4e + 4\ K$ is the reciprocal of the Boltzmann's constant, $T(x_v(\tau), y_v(\tau), z_v(\tau))$ is the temperature in a point $(x_v(t), y_v(t), z_v(t))$ along the trajectory $v$, and $\alpha$ being chosen so that $P_{v0}(t = 6h, T = 0) = 0.01$. The exponent is divided by 24 because $t$ is measured in hours. We made this choice because we wanted to mimic a progressive loss of spores due to dilution and dry deposition, assuming that viable spores become negligible after 6 hours.

To obtain the probabilities of deposition $P_v$, we corrected $P_{v0}$ to remove those parts of the trajectories whose altitude is higher than the planetary boundary layer.

Eventually, we performed a last correction to account for the fact that spore deposition is directly proportional to the time spent by the air mass over the crops. Let us consider $l_h$ the Euclidean distance between two consequent points $(x_h, y_h)$ and $(x_{h+1}, y_{h+1})$ of a trajectory, where $h$ indicates the time in hours. This proportionality means that, the longer the segment $l_h$, the lower the probability of spore deposition in a point of that segment. Therefore, we computed $P_{v,t}$ as:

$$P_{v,t} = P_{v0,t}(1 - \frac{l_h - \min_{\forall h \in 0,1...6} l_h}{\max_{\forall h \in 0,1...6} l_h}) \tag{6}$$

In this, way, in the shortest segment ($\min_{\forall h \in 0,1...6} l_h$) the value of $P_{v,t}$ is equal to that of $P_{v0,t}$, since this latter is multiplied by 1, while the longest ($\max_{\forall h \in 0,1...6} l_h$) is multiplied for the ratio between the shortest and the longest segment ($\min_{\forall h \in 0,1...6} l_h / \max_{\forall h \in 0,1...6} l_h$), which is less than (or equal) to 1.

#### 1.6 Connectivity matrix

Once we have obtained matrices $W_t$, which consider all trajectories $v$ that started in $t$, we first performed a temporal aggregation on a daily basis $d$ (where $t$ is intended as a discrete measure of time, while $d$ refers to a specific day of the year) and secondly corrected to account for neighbouring units:

1. The temporal aggregation consists of averaging all the connectivity matrices $W_t$ obtained on the same day of the year. For instance $W'_d$ on the 15$^{\text{th}}$ of May (i.e., $W_{15-05}$) contains the element wise average of all the $11 \times 4 = 44$ corresponding $W_t$ computed on the 15$^{\text{th}}$ of May 2008, 2009,... 2018, computed from Lagrangian simulations launched four times a day;

2. The correction for neighbouring units accounts for geographic proximity. We consider that two bordering units may reciprocally infect even when we have not found any connection among them. We first considered all couples of 8-neighbouring units $i$ and $j$, then computed the average connectivity $\tilde{w}$ among all $W'_{d,ij}$. We then corrected matrices $W'$ as follow:

$$W_{d,ij} = max(W'_{d,ij}, \tilde{w}) \tag{7}$$

$W_d$ is the matrix used in the metacommunity model. We calculated also the annual matrix $W$, which is the average of all $W_t$ during all ripening season in 2008-2019, where a filter has been added to consider only rainy events in the arrival unit, which can lead to wet deposition (which means that each element of $W$ is determined by $W_{ij} = (1/T)\sum_t^T (r_{t,j} > 0)W_{t,ij}$)

#### 1.7 Determination of the prior probability function

We set a tentative prior distribution of $\boldsymbol{\theta}$, which we iteratively refined via simulations (see Godding et al., 2022).

We estimated the shape of the prior distribution of parameter $\theta_O$, which weights the probability of overwintering (i.e., the probability of having an inoculum at the beginning of the next season because of the presence of mummies from the previous ripening reason), by considering the observation dataset. Such a probability may be approximated by the ratio between $i)$ the occurrences of a "strong" disease incidence in year $y+1$ in early or mid-early cultivar after a "strong" incidence in year $y$ in the same location over $ii)$ the occurrences of a high incidence in year $y$. This ratio is equal to 1/3 (2 occurrences in the observation dataset over 6). So, we started with the hypothesis that the expected value $\boldsymbol{E}[.]$ of the random variable corresponding to presence of local inoculum at the beginning of the season ($\boldsymbol{E}[E_y(t_0) = 0.27\ fruits/m^2]$, hereafter $P_o$) is defined as follows:

$$P_o = \boldsymbol{E}[1 - (1 - \tilde{P}_o)(e^{-\theta_O M_{t_f,y-1}})] = 1/3 \tag{8}$$

Where $\tilde{P}_o$ represents the presence of spores due to external causes, while $1 - e^{-\theta_O M_{i,t_f,y-1}}$ expresses the probability of overwintering, with the density of mummies at harvest time of the precedent year $M_{t_f,y-1}$ weighted by parameter $\theta_O$. We estimated $\tilde{P}_o = 0.2$ directly from the observations dataset as the frequency of "weak" disease severities followed by a "strong" disease severity in early and mid-early cultivars of the following year in the same place (2 occurrences in the observation dataset over 10), under the simplification assumption that, due to shorter time, inoculum from overwintered mummies could have been the main cause of infection in early and mid-early cultivars.

Considering $M_{t_f,y-1} = M_0 = 10\ fruits/m^2$ as a strong infection, we obtained:

$$\boldsymbol{E}[\theta_O] = \frac{1}{M_0} log(\frac{1-\tilde{P}_o}{1-P_o}) = 0.018 \tag{9}$$

We than considered $\theta_O$ as extracted from an exponential variable with average $1/\boldsymbol{E}[\theta_O]$, which is $\approx 54.8$. The exponential variable allows us to obtain $i$) positive values, $ii$) scattered around $\boldsymbol{E}[\theta_O]$, $iii$) allowing us to explore also few very large values, with a parameter only ($1/\boldsymbol{E}[\theta_O]$). While launching preliminary ABC algorithms with a reduced number of simulations, we progressively brought the value of $1/\boldsymbol{E}[\theta_O]$ up to 81, since this increased the number of accepted parameter sets.

We have several simulations spanning several order of magnitudes to find the interval in which $\theta_E$ describes both $i$) almost no new introductions and $ii$) almost the maximum of introduction each year. Eventually, the prior distribution of $\theta_E$ was set as an exponential of a uniform distribution between $-8log(10)$ and $-5log(10)$.

Parameter $\theta_L$ is to be considered as the losses threshold over which a disease incidence is considered as “strong”. From consistency with how the observation dataset was built, we assumed that this parameter should be placed between 0.2 and 0.4. We therefore considered a lognormal distribution, with parameters $\mu_L = -1.26$ and $\sigma_L = 0.21$, for which 0.2 and 0.4 represent the $5^{\text{th}}$ and the $95^{\text{th}}$ percentiles, respectively. We selected a lognormal distribution for similar reasons that brought us to choose an exponential variable for overwintering, but with enough indications to guess two parameters instead of one. While launching preliminary ABC algorithms with a reduced number of simulations, we updated the values of $\mu_L$ and $\sigma_L$ respectively to 1.2 and 0.15.

#### 1.8 Training and stratified cross validation

In our training, we ran 200,000 simulations, spanning over 1996-2021, under the assumption that initial inoculum is randomly present in each unit with probability $\tilde{P}_o$.

We split the observations $\Omega = 5$ times into a training set $\phi$ (containing $1 - 1/\Omega = 80\%$ of the observations, over which ABC was applied) and a testing set $\psi$, representative of the observations structure.

We formed $\Omega = 5$ testing set so that each one satisfies all the following criteria: $i$) it includes $1/\Omega = 20\%$ of observation dataset; $ii$) it includes all the locations of the observation dataset (Fig.

3 of the main text) and cultivars; *iii*) where possible (given the reduced degrees of freedom due to criteria *i* and *ii*), locations and cultivars are represented proportionally to the complete dataset .

We relaxed a fourth criteria, i.e. *iv*) disease incidences represented proportionally to the complete dataset, since it was not compatible with *i*)-*iii*); we considered only testing sets in which proportions the ratio of "weak" and "strong" are not too distant from the proportions in the complete dataset: they should not exceed 60% and 45% of the testing set, respectively.

Criteria *i* to *iv* allow us to "stratify" the cross validation, i.e. they assure that each testing set is a good representation of the whole dataset. However, this "stratification" has the drawback that some observations repeat systematically in the different testing sets. If this repetition becomes important, to the limit where all testing sets contain the same observation, this can represent a problem, since it means that the model validation is performed against only one the same testing set, which reduces the model's confidence. Therefore, we computed the grade of similarity among the testing sets (i.e., the ratio of observations which are identical for two sets), which is between 18.5% and 37.5%. We considered it acceptable since it never represent the largest part (its grade of similarity being always strictly lower than 50%).

In order to choose the 100 more frequent accepted sets, we weighted frequencies by the corresponding $\kappa_{\psi,\omega}$ in split $\omega$

#### 1.9 Null model: isotropic kernel

We computed matrix $\mathbf{U}$ by computing each element $u_{ij}$ connecting $i$ and $j$ (which are found $d$ units away), as the average of all $w_{mn}$, where $m$ and $n$ are all the couples of nodes which are $d$ units away. A representation of the kernel is given in Fig. SI7a.

The Monte Carlo analysis is performed this way this way:

1. the posterior distribution $\boldsymbol{\theta^U} = (\theta_E^U, \theta_O^U, \theta_L^U)$ is cumputed via ABC with 200.simulations;

2. Named $\kappa_w$ and $\kappa_u$ the vectors of $\kappa$ computed with the prior distribution of the parameters, we computed $D_o = \overline{\kappa}_w - \overline{\kappa}_u$;

3. To check if $D_o$ is larger than it would be by chance, we created $20,000$ vectors $\hat{\kappa}_w$ and $\hat{\kappa}_u$ by reshuffling elements of $\kappa_w$ and $\kappa_u$, and computed $\hat{D}_i$ as $\overline{\hat{\kappa}}_{w,i} - \overline{\hat{\kappa}}_{u,i}$;

4. We compared the distribution of $\hat{D}_i$ with respect to $D_o$.

##### 1.10 Definition of "vulnerability" and "dangerousness"

Given any unit $j$ ($j = 1, 2, \ldots N = 755$) characterised by its cultivated area $Ac_j$, a time horizon defined by $y = 10\ years$, a stochastic iteration $\lambda$ of the model ($\lambda = 1, 2, \ldots, \Lambda = 100$), local losses $L_{ij,y}$ (defined by Eq. 5 in the main text) due to an inoculated unit $i$ ($i = 1, 2, \ldots N = 755$), vulnerability $v_j$ and dangerousness $d_i$ are defined by Eq.s 10 and 11:

$$v_j = \frac{1}{\Lambda(N-1)} \sum_{\lambda, i \neq j} L_{ij,y,\lambda} \tag{10}$$

$$d_i = \frac{1}{\Lambda \sum_{j \neq i} Ac_j} \sum_{\lambda, j \neq i} L_{ij,y,\lambda} Ac_j \tag{11}$$

Note that, since vulnerability computes the local losses, it is normalized by the number of cells reduced by one ($N - 1$), while in dangerousness one wants to weight losses according to the cultivated surface of the other cells ($\sum_{j \neq i} Ac_j$).

In these equations we imposed $j \neq i$ so that to exclude the contribution of the first inoculated units. Otherwise, these two indices would be artificially biased toward units with large $Ac$.

### 2 Supplementary Results

#### 2.1 Domain extraction

Peach surface by NUTS2 region is summarized in Tab. 1. The final values of peach orchards surface per unit is reported in Fig. SI1e.

#### 2.2 Spatial distribution of cultivars

The probability for each cultivar (early, mid-early, mid-late, late) of being chosen in the meta-community model is represented in Fig. SI1a-d. The sum of the probabilities for each cultivar is 100%.

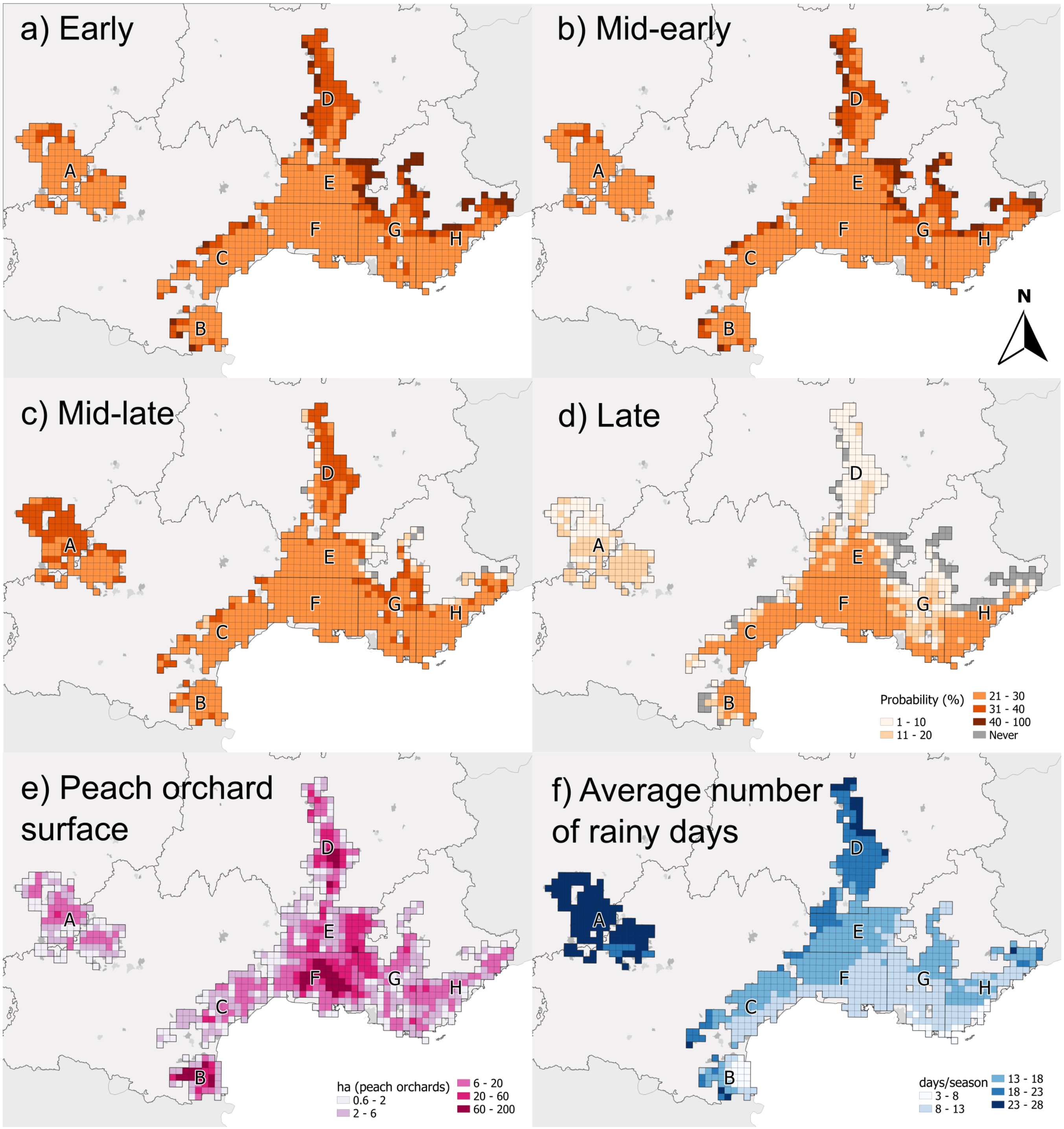


**Figure 1:** a - d) Probability of being chosen for each cultivar e) peach orchard surface by unit in *ha*; f) average number of rainy days during each ripening season. Regions of Fig. 2b of the main text are reported.

**Table 1:** Amount of peach surface by French NUTS2 region in 2017.

| NUTS2 | Peach surface (ha) |
|---|---|
| FR1 (Île-de-France) | 0 |
| FR2 (Champagne-Ardenne, Picardy, Upper Normandy, Centre, Lower Normandy, Burgundy) | 24.8 |
| FR3 (Nord-Pas-de-Calais) | 0 |
| FR4 (Lorraine, Alsace, Franche-Comté) | 12.5 |
| FR5 (Pays de la Loire, Brittany, Poitou-Charentes) | 27.6 |
| FR6 (Aquitaine, Midi-Pyrénées, Limousin) | 629.2 |
| FR7 (Rhône-Alpes, Auvergne) | 2,007.8 |
| FR8 (Languedoc-Roussillon, Provence-Alpes-Côte d'Azur, Corsica) | 7,879.0 |

#### 2.3 Average connectivity matrix

The value of $\tilde{w}$ is 0.053. A detailed version of the average air-masses driven connectivity matrix $W$ (depicted as network in the main text, Fig. 2b) is represented in Fig. SI2.

#### 2.4 ABC, stratified cross validation

The results of the ABC applied over the $\Omega = 5$ stratified splitting of the observation dataset is reported in Tab. 2. The overall 100 parameters sets are represented as marginalized posterior function in Fig. SI3. To facilitate interpretability of the accepted parameters $\theta_E$ and $\theta_W$, the "average annual occurrences of local primary inocula" and the "average annual introductions of external inocula" are represented in Fig. SI4.

**Table 2:** Parameter's performances in training and testing.

| Repetition ($\omega$) | 1 | 2 | 3 | 4 | 5 |
|---|---|---|---|---|---|
| Cohen's $\kappa_\phi$ (training set) | 0.496 | 0.497 | 0.502 | 0.492 | 0.502 |
| Cohen's $\kappa_\psi$ (testing set) | 0.046 | 0.086 | 0.289 | 0.409 | 0.161 |
| Accepted parameter set size | 67 | 34 | 19 | 7 | 18 |

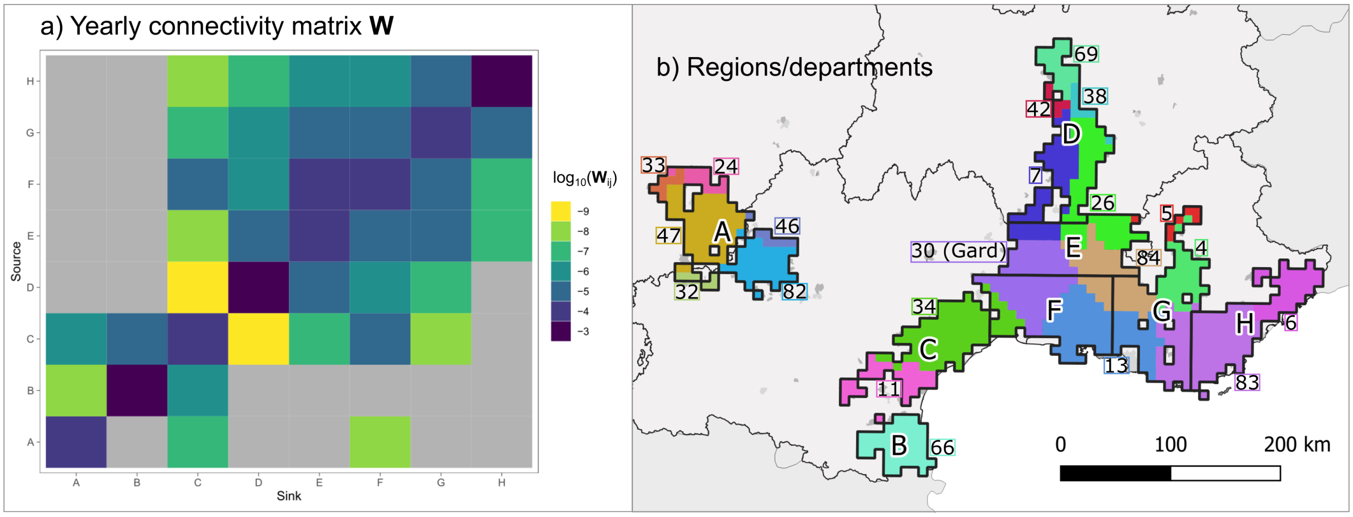


**Figure 2:** a) Average yearly connectivity matrix $W$, represented grouping units according to regions in panel b), where, in addition to regions, also French administrative departments are mapped with different colours. The colour scale of panel a) is built to represent the $log_{10}$ of each element $W_{ij}$. In panel b), department 30 (Gard, in violet in the center) is the one where *M. fructicola* has been detected first in 2001 (EPPO, 2023).

#### 2.5 Model accuracy

The average correct classification rate (Fielding and Bell, 1997), i.e. the frequency of correspondences (true positives and true negatives) between the observation dataset and the model's output, grouped by observation, is represented in Fig. SI5.

#### 2.6 The null model

Depending on the cross validation splitting $\omega$, the size of the accepted parameter sets varies from 10 to 38 (average = 18). Values of $\kappa_\phi$ have an average of 0.5 (min = 0.495; max = 0.510), while $\kappa_\psi$ has an average value of 0.11 (min = 0.07, max = 0.16). The size of the accepted parameter set is 68.

Accuracy of the parameterization of the null model per location, year and variety is reported in Fig. SI6, while accepted parameters are reported in Tab. SI3. Matrix $U$ used in the null model is represented in in Fig. SI7a.

Results of Monte Carlo analysis are reported in Fig. SI7b. The statistical distribution of the

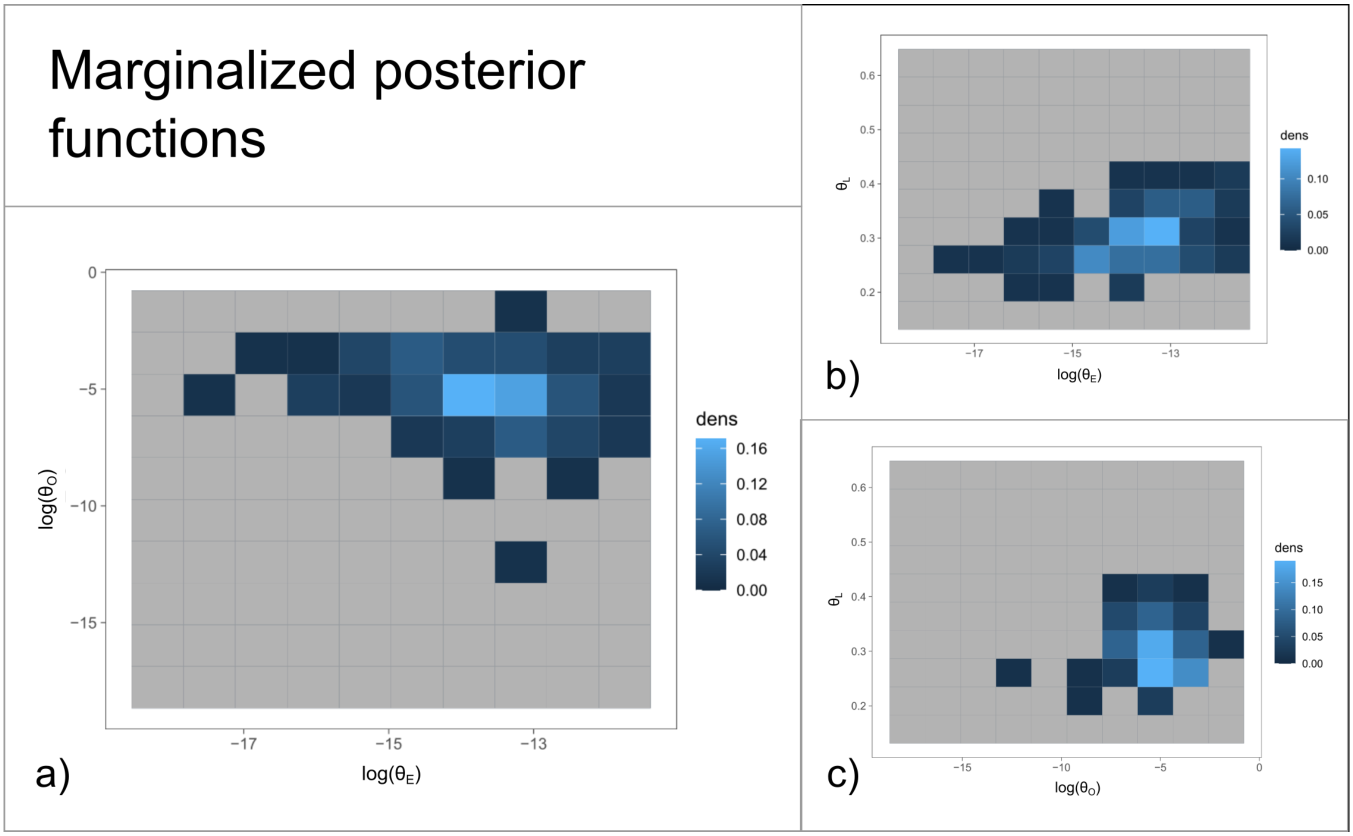


**Figure 3:** Panels a) to c) show the marginal distribution of the posterior function for all possible a couple of the three parameters $\theta$. Lighter the colour, higher the probability. Note that $\theta_L$'s axis is linear, while the others' are logarithmic. In a) the posterior distribution is normalized over $\theta_E$ and $\theta_O$; this panel is larger since it represents the parameters of the main modules of the metacommunity model (i.e., external inoculum, Fig. 1d in the main text, and interannual persistence of inoculum, Fig. 1e in the main text). In b) the posterior distribution is marginalized over $\theta_E$ and $\theta_L$, while in c) the posterior distribution is marginalized over $\theta_O$ and $\theta_L$.

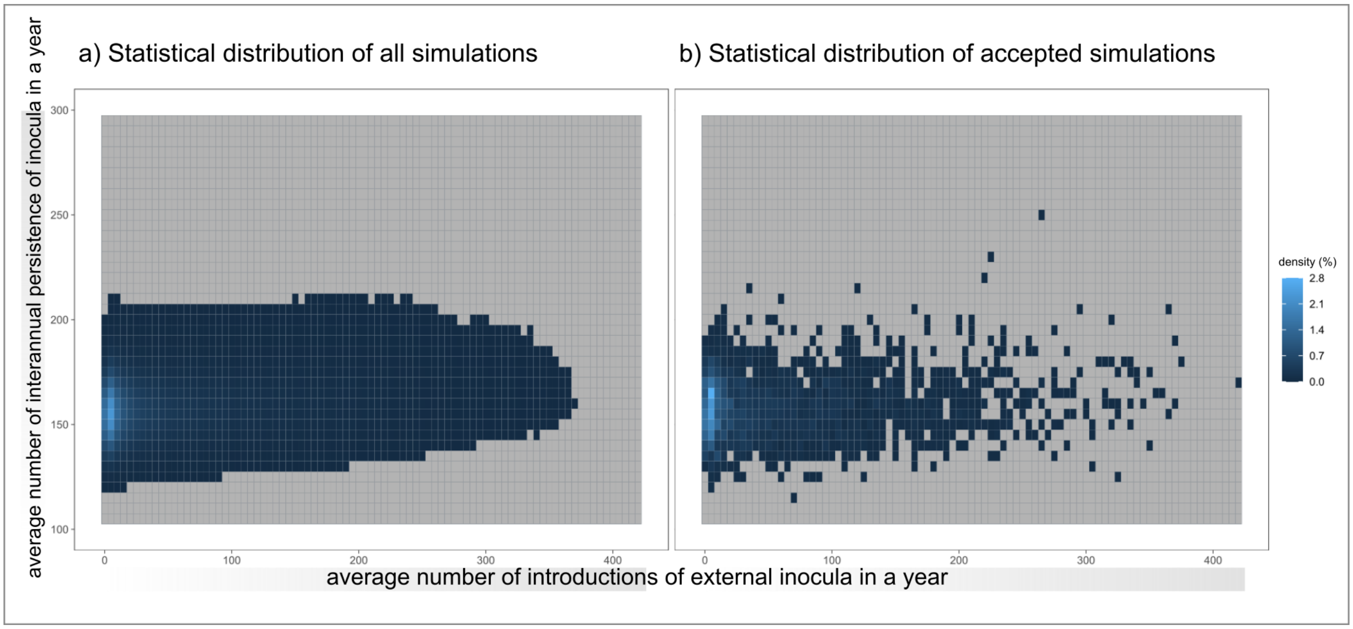


**Figure 4:** Statistical distribution of "average number of introductions of external inocula in a year" and "average number of interannual persistence of inocula in a year" in a) all the simulations and b) the simulations of the ensemble of the accepted parameter sets. Density values $< 0.1\%$ have been omitted.

difference $\hat{D}_i = \hat{\kappa}_{w,i} - \hat{\kappa}_{u,i}$, where $\hat{\kappa}_{u,i}$ and $\hat{\kappa}_{u,i}$ ($i = 1, \ldots, 20,000$) contain randomly reshuffled elements of $\kappa_w$ and $\kappa_u$, is always lower than the observed difference $D_o = \kappa_w - \kappa_u$. There is no reshuffled distances $\hat{D}_i$ (average value $= -3.01 \times 10^{-6}$; interquartile range $= -2.11 \times 10^{-4}$ to $2.04 \times 10^{-4}$) which is larger than the observed distance between the mean performance of the full model $D_o$ $9.50 \times 10^{-3}$.

**Table 3:** Parameter's performances in training and testing of the null model.

| Repetition ($\omega$) | 1 | 2 | 3 | 4 | 5 |
|---|---|---|---|---|---|
| Cohen's $\kappa_\phi$ (training set) | 0.497 | 0.510 | 0.496 | 0.498 | 0.495 |
| Cohen's $\kappa_\psi$ (testing set) | 0.076 | 0.068 | 0.155 | 0.113 | 0.152 |
| Accepted parameter set size | 38 | 12 | 15 | 10 | 13 |

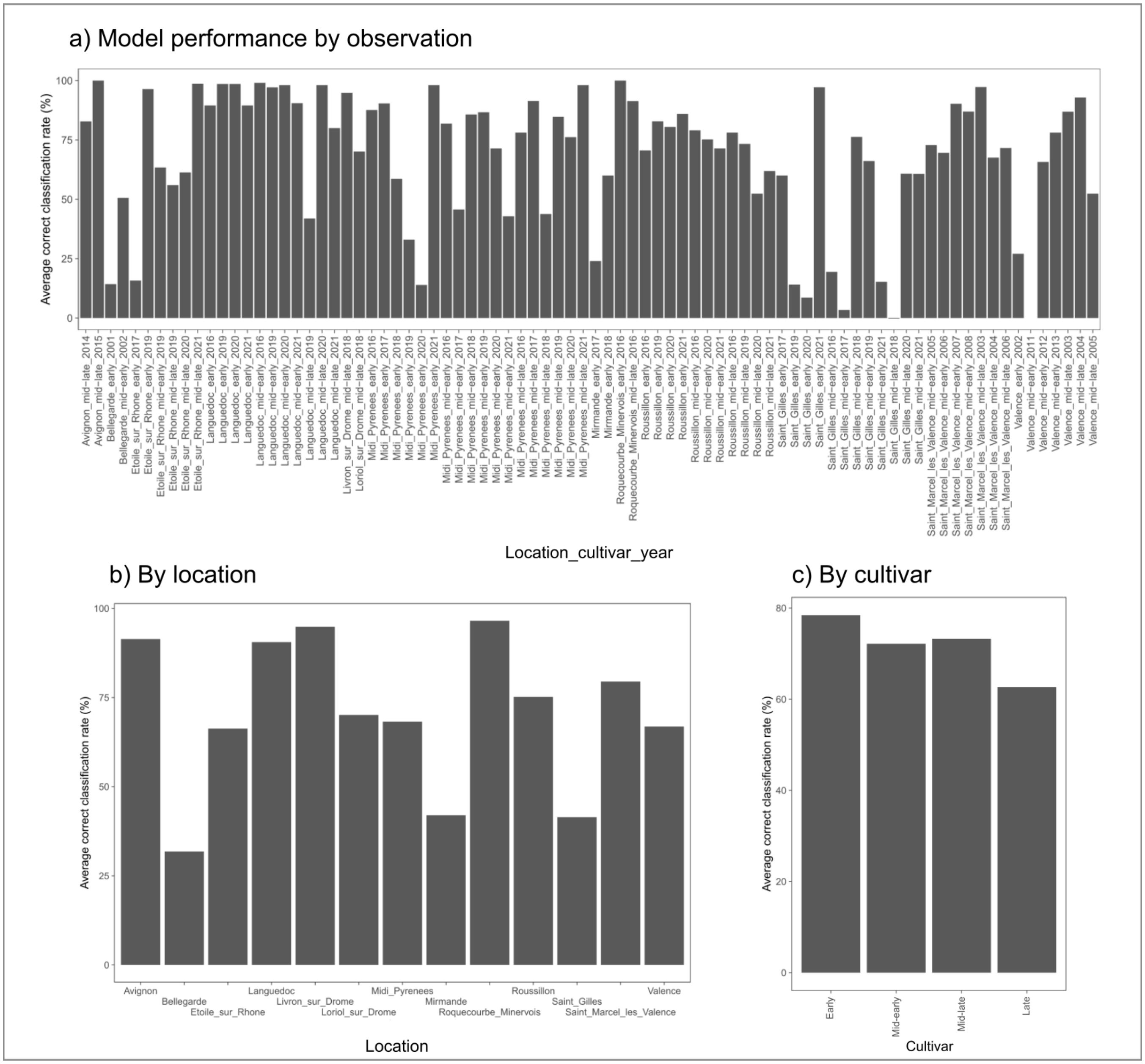


**Figure 5:** Average correct classification rate (Fielding and Bell, 1997), expressed as %, a) by observation b) by location and c) by cultivar. Note that panel a) represents the same rates as in Fig. 3 in the main text.

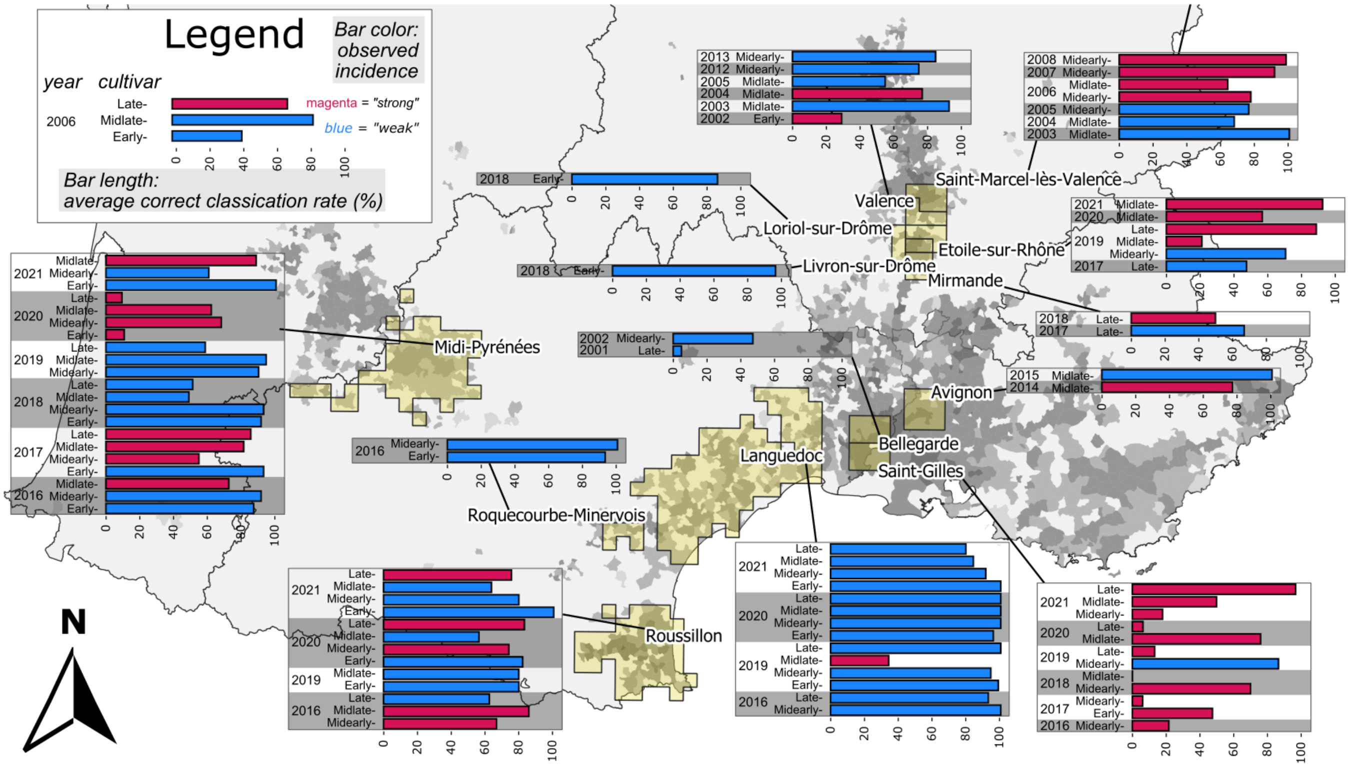


**Figure 6:** Summary of the performances of the null model throughout the domain. The length of the bars represented the average correct classification rate of the null model with the accepted parameter set. The legend is equivalent of that of Fig. 3 of the main text.

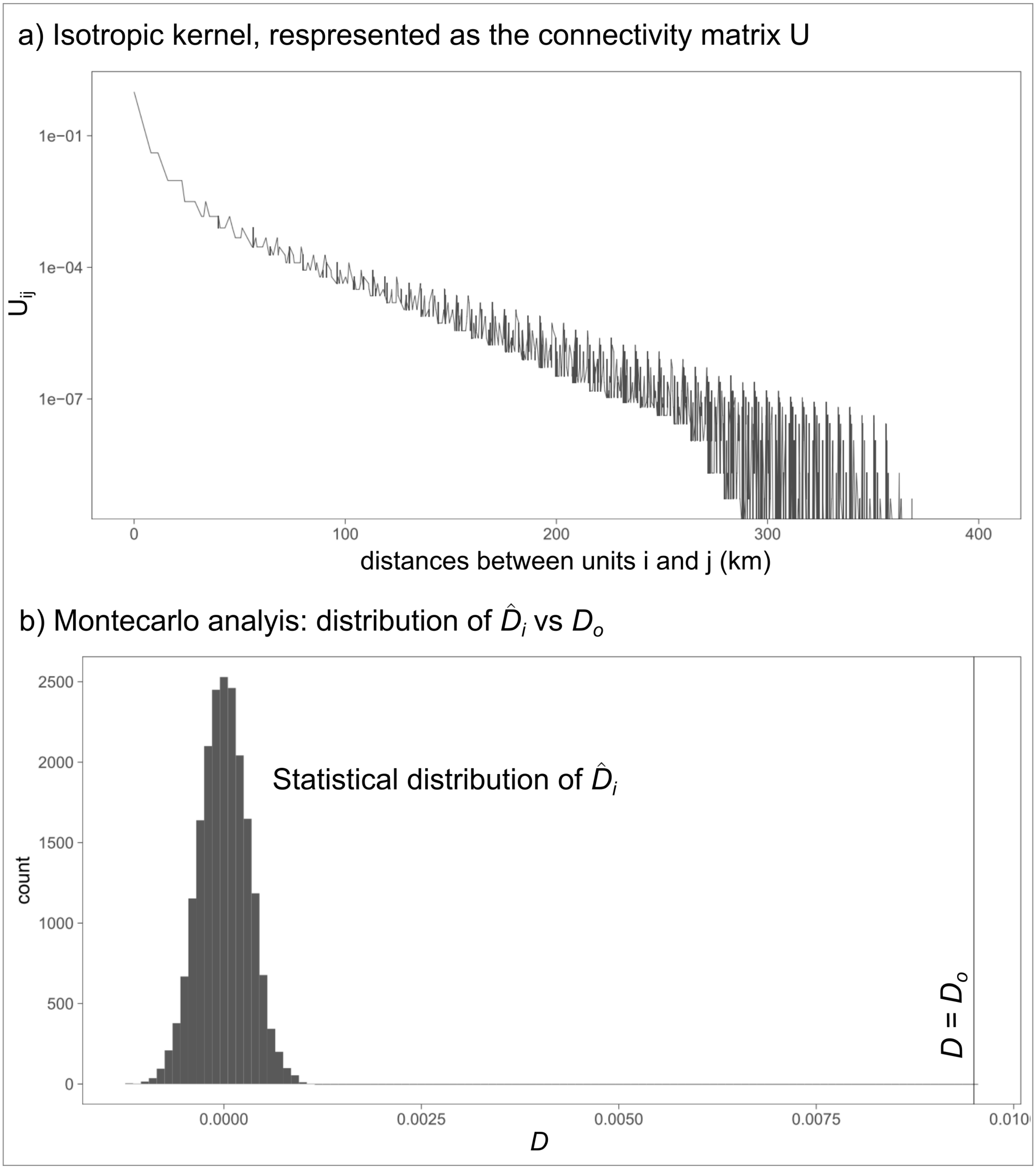


**Figure 7:** Null model: a) representation of each element of the matrix $U_{ij}$ as a function of the distance (in km) between units $i$ and $j$; b) representation of the Montecarlo analysis. The statistical distribution of the difference $\hat{D}_i = \hat{\kappa}_{w,i} - \hat{\kappa}_{u,i}$, where $\hat{\kappa}_{u,i}$ and $\hat{\kappa}_{u,i}$ ($i = 1, \ldots, 20,000$) contain randomly reshuffled elements of $\kappa_w$ and $\kappa_u$, is compared with the observed difference $D_o = \kappa_w - \kappa_u$. No element $\hat{D}_i$ is greater than $D_o$.

QGIS Development Team, QGIS Geographic Information System. QGIS Association, 2022.

Peach and apricot trees - Area by density classes and group of varieties (area in ha). shorturl.at/afNVY, 2021; Accessed: 2022-07-07.

Bevacqua, D.; Vanalli, C.; Casagrandi, R.; Gatto, M. A climate-driven compartmental model for fungal diseases in fruit orchards: The impacts of climate change on a brown rot-peach system. *Agricultural and Forest Meteorology* **2023**, *332*, 109293.

Vanalli, C.; Casagrandi, R.; Gatto, M.; Bevacqua, D. Shifts in the thermal niche of fruit trees under climate change: The case of peach cultivation in France. *Agricultural and Forest Meteorology* **2021**, *300*.

Delannoy, D.; Maury, O.; Décome, J. CLIMATIK : système d'information pour les données du réseau agroclimatique INRAE. 2022; https://doi.org/10.57745/AJNXEN.

Caubel, J.; García de Cortázar-Atauri, I.; Launay, M.; de Noblet-Ducoudré, N.; Huard, F. F.; Bertuzzi, P. P.; Graux, A.-I. Broadening the scope for ecoclimatic indicators to assess crop climate suitability according to ecophysiological, technical and quality criteria. *Agricultural and Forest Meteorology* **2015**, *207*, 94 – 106.

Godding, D.; Stutt, R. O. J. H.; Alicai, T.; Okao-okuja, G.; Gilligan, C. A. Developing a predictive model for an emerging epidemic on cassava in sub-Saharan Africa. *bioRxiv* **2022**, 1–24.

EPPO, Monilinia fructicola. EPPO datasheets on pests recommended for regulation. https://gd.eppo.int, 2023; Accessed: 2023-05-09.

Fielding, A. H.; Bell, J. F. A review of methods for the assessment of prediction errors in conservation presence/absence models. *Environmental Conservation* **1997**, *24*, 38–49.
